# Physical factors contributing to regulation of bacterial surface motility

**DOI:** 10.1101/719245

**Authors:** Ben Rhodeland, Kentaro Hoeger, Tristan Ursell

## Abstract

Microbes routinely face the challenge of acquiring territory and resources on wet surfaces. Cells move in large groups inside thin, surface-bound water layers, often achieving speeds of 30 µm/s within this environment, where viscous forces dominate over inertial forces (low Reynolds number). The canonical Gram-positive bacterium *Bacillus subtilis* is a model organism for the study of collective migration over surfaces with groups exhibiting motility on length scales three orders of magnitude larger than themselves within a few doubling times. Genetic and chemical studies clearly show that the secretion of endogenous surfactants and availability of free surface water are required for this fast group motility. Here we show that: (i) water availability is a sensitive control parameter modulating an abiotic jamming-like transition that determines whether the group remains fluidized and therefore collectively motile, (ii) groups self-organize into discrete layers as they travel, (iii) group motility does not require proliferation, rather groups are pulled from the front, and (iv) flow within expanding groups is capable of moving material from the parent colony into the expanding tip of a cellular dendrite with implications for expansion into regions of varying nutrient content. Together, these findings illuminate the physical structure of surface-motile groups and demonstrate that physical properties, like cellular packing fraction and flow, regulate motion from the scale of individual cells up to length scales of centimeters.

## Introduction

In their search for resources microbes contend with physically distinct environments ranging from soft surfaces to bulk Newtonian fluids and complex fluids like mucus. In bulk fluid environments, canonical microbes like *Escherichia coli* (1) and *Bacillus subtilis* (2, 3) ascend favorable chemical gradients via run-and-tumble chemotaxis (4). This mechanism of gradient ascent requires both flagellar-mediated motility and a complex system of phosphorylation-memory and chemical sensors on the bacterial surface, that together regulate the run-tumble transition frequency (5). In contrast, bacterial surface motility has different requirements and inputs, and different species have evolved distinct modalities of surface motion. For instance, in the predatory species *Myxococcus xanthus*, individual cells move in back-and-forth motions that employ two distinct sets of protein machinery for ‘twitching’ and ‘gliding’ motility, and cells assemble into larger motile groups that traverse surfaces as monolayers (6, 7). Many other species, including the opportunistic pathogens *Serratia marcescens* (8, 9) and *Proteus mirabilis* (10, 11), also form large groups of motile cells that are capable of rapidly expanding over surfaces, in some cases even against bulk fluid flow (12). Similarly, when present in sufficient numbers *Paenibacillus dendritiformis* form intricate fractal-like patterns on soft agar surfaces in response to lateral chemical gradients (13, 14). Even baker’s yeast have been observed to exhibit group movement over fluid surfaces (15). Other species exhibit surface motility in response to non-chemical fields; for instance the cyanobacterium S*ynechocystis* is phototactic, responding to incident light by asymmetrically extending and retracting pili from its surface to create a biased random walk toward a light source (16, 17). Crucial to its motion, *Synechocystis* modifies the local surface environment by secreting exopolysaccharides, and only when enough cells have participated in such surface modification can the group move toward the light source. These examples demonstrate that in response to various gradients, microbes have evolved distinct sensing capabilities and modalities of motion to acquire resources and respond to selective pressures on surfaces. Despite their differences, surface motility in all of these species (and even in abiotic systems (18)) appears to be a collective phenomenon, requiring the motion of and/or surface modification by large numbers of cells (19, 20). Uncovering biochemical and genetic factors that regulate motion is crucial for understanding how each species executes surface motility, but those factors are only part of the full picture. Physical forces that produce, regulate, and guide microbial group motion on surfaces may be relevant across species and contexts, and are thus integral to our understanding of microbial ecology in natural environments and will expand the suite of design tools for engineering microbial systems.

Modeling plays an important role in these systems because it has the potential to connect experimental observations with physical forces and regulators of motion (21–29). Different models posit qualitatively distinct physical mechanisms for group motility, with basal assumptions motivated by species-specific attributes. In *P. dendritiformis* growth and spreading of cells in dendritic patterns are modeled as a diffusion-limited conversion of nutrients into biomass at the growing tips of the dendrite (13, 30). Cellular motion is thought to rely on chemotaxis that follows nutrient gradients that are created by metabolic consumption of the colonies themselves (22, 30, 31). Together these effects recapitulate many of the classes of bacterial surface patterning observed in experiments (26, 32–36). However, within a dense and active (e.g. swarming) group whose velocity correlations rapidly decay on the length scale of a few cells (21), it is unclear how individuals could effectively modulate their tumble frequency and accumulate sufficiently persistent runs to deliberately bias their random walks and hence execute run-and-tumble chemotaxis. Indeed, previous work in *B. subtilis* (37–40) (and *Pseudomonas aeruginosa* (41), *Salmonella enterica* (42), and *Escherichia coli* (43)) shows that neither chemotaxis nor motility of individuals are necessary for rapid and outward-directed surface motility. Other models of surface motility (e.g. swarming or sliding) in *B. subtilis* or *P. aeruginosa* focus on the role of surfactants (23, 44) and/or osmotic potential (24), both of which appear to be crucial for rapid surface motility in those species (37, 45–47).

While cell density is clearly an important factor in motion, in so much as many cells are required for collective movement, and genetics implicates endogenous surfactant production as a physical requirement for collective motion, it remains unclear how density, material flow, and dendrite structure contribute to regulation of this ubiquitous behavior. In this work we combined high resolution video and time-lapse microscopy with computational image processing to clarify the relative contributions of density, flow and structure by examining the surface motility of the extensively-studied Gram-positive bacterium *B. subitilis* (5, 24, 48, 49). Within a few hours of deposition, a small central inoculum of wild-type cells can rapidly colonize the entire surface of a wet 10 cm agar plate via apparent swarming motility (50). Such group motility over soft surfaces has been shown to depend on the secretion of ‘surfactin’, a bacterially produced surfactant and wetting agent (22, 23, 25, 37, 51–53). Functional knockouts for surfactin production (Δ*srf*) result in a phenotype where individual cells are still motile and chemotactic in bulk fluid, but bacterial groups cannot move across surfaces (37, 40, 54). Localized secretion of surfactin is thought to generate a gradient in surface tension, and thus produce collective motion via the Marangoni force (23, 55, 56).

Building on that biophysical picture, we found evidence that the movement of *B. subtilis* over soft surfaces was regulated by cell density, whereby groups of cells are subject to a jamming-fluidization transition (57) that correlates with packing fraction (and thus water availability). These results complement findings that the viscosity of bacterial suspensions depends on cell concentration, shear rate, and level of motility (58, 59). Further, we show that such groups operate in two distinct modalities of motion: groups can move without growth as ‘islands’ that translocate independent of the parent colony, or dendrites can extend with large-scale flow of material from the parent colony to the dendrite tip. We used a chemotropic assay to show that when connected to a nutrient-rich parent colony, extending dendrites can venture into nutrient-barren regions, while colonies that originate in nutrient-barren regions are not collectively motile. These data suggest that, in addition to genetically regulated production of surfactants, multiple physical and abiotic factors regulate motion independent of the chemotactic or motile abilities of the constituent cells, and that groups move differently when presented with anisotropic nutrient environments. When combined with previous work, these data suggest a model in which individual motility and subsequent swarming are mechanisms that maintain a fluidized state on which surface-tension gradient forces (Marangoni forces) and osmotically driven colony hydration can act to precipitate group motion over a surface, and that a jamming-like transition, like those found in other macroscopic granular systems (57, 60–65), may be a key regulator of whether bacterial groups are able to move.

## Results

*Bacillus subtilis* is a model motile Gram-positive bacterium, capable of sensing and responding to its chemical environment in bulk fluid via run-and-tumble chemotaxis (2, 5, 66). On wet surfaces wild-type *B. subtilis* rapidly move out from a central inoculum (67), apparently via collective swarming motility (50) that requires secretion of the endogenous bio-surfactant ‘surfactin’. Mutants that lack the ability to produce surfactin (Δ*srf*) do not exhibit collective motility from their central inoculum (37) (and Movie S1). We inoculated small, dense (OD ∼ 10 - 20) droplets of wild-type *B. subtilis* on soft (∼0.5% w/v agarose) nutrient-rich surfaces. After a brief quiescent phase, groups of cells rapidly expanded over the surface in a dendritic pattern that reached the edge of the plate (∼5 cm of travel) in less than 6 hours (Movie S2). Dendrites robustly moved outward away from the original point of inoculation into fresh territory at an average group motility rate of ∼5 µm/s and up to ∼15 µm/s (SI Fig. 1). Cells within the dendrites were highly motile, exhibiting swarming motility (37) with individual cells moving at rates up to ∼30 µm/s in highly circuitous paths. Consistent with previous work (37, 38), we confirmed that mutants lacking the ability to run (tumble-only, Δ*CheY*), lacking the ability to tumble (run-only, Δ*CheB*), and mutants lacking flagella (Δ*hag*) were all able to exhibit rapid surface motility, albeit with some differences in dendrite speed and spreading pattern (SI Fig. 2, Movies S3 - S5).

In our early observations we noted transitions in the motile state of groups, wherein cells within a *contiguous region* appeared to be either highly motile and moving as a group or immobilized on both the individual and group levels. These distinct states of motion were frequently observed at the same time, and cellular groups were observed transitioning between these states on the time scale of a minute, too short to be accounted for by phenotypic changes.

### Movement is regulated by a jamming-like transition

While surface motility of *B. subtilis* is robust to genetic manipulations of chemotaxis and motility, it was very sensitive to gel stiffness as measured by agarose concentration (37). Below ∼0.4% agarose by weight the gel is sufficiently porous that bacteria can penetrate and swim within it, akin to canonical swim plates (50, 68). Above ∼0.7% agarose, limited water availability hinders surface motility (50), leading to expansion across plates through the replication of sessile cells at the leading edge of growth, as seen in *P. dentritiformis* (dendrite-like) or *E. coli* (growing circular colonies), both of which expand at much slower rates (31, 69, 70). Thus, across a relatively narrow range of agarose percentages, groups of *B. subtilis* exhibit three qualitatively distinct behaviors: swimming through agarose (below ∼0.4%), rapid surface motility (between ∼0.4% and ∼0.7%), and slow growth by replication (above ∼0.7%). Why does the transition from rapid surface motility to slow proliferative growth occur over such a narrow range of agarose stiffness, and correspondingly, water availability?

Many previous studies of *B. subtilis* characterized spreading and attendant morphological behaviors at the length-scale of colonies (mm to cm) and on the timescale of bacterial replication, imaging colony morphology and spreading with minutes or hours between frames (26, 32, 37, 50). These excellent studies revealed much of what is known about spreading phenotypes, their genetic mechanisms, and biochemical correlates. We performed high temporal and spatial resolution imaging of spreading colonies to illuminate processes that potentiate spreading. We captured images at 30 or 60 frames per second with spatial resolution of 5 µm/pixel or 15 µm/pixel, respectively, both of which enabled us to see intensity variations produced by the movements of cells within the swarm. We wrote a custom image analysis script that measured a scalar correlate of motion as a function of position through time. Briefly, the algorithm measures the mean absolute value of local intensity fluctuations at a position across a set of *N* (usually 5 – 7) frames and thus reports on the level of motile activity at each position through time (see Methods). With this ‘activity’ filter, we were able to visualize on the scale of 10s to 100s of microns which parts of the colony were actively motile and in which parts cells were stationary (Fig. 1A).

**Figure 1.**
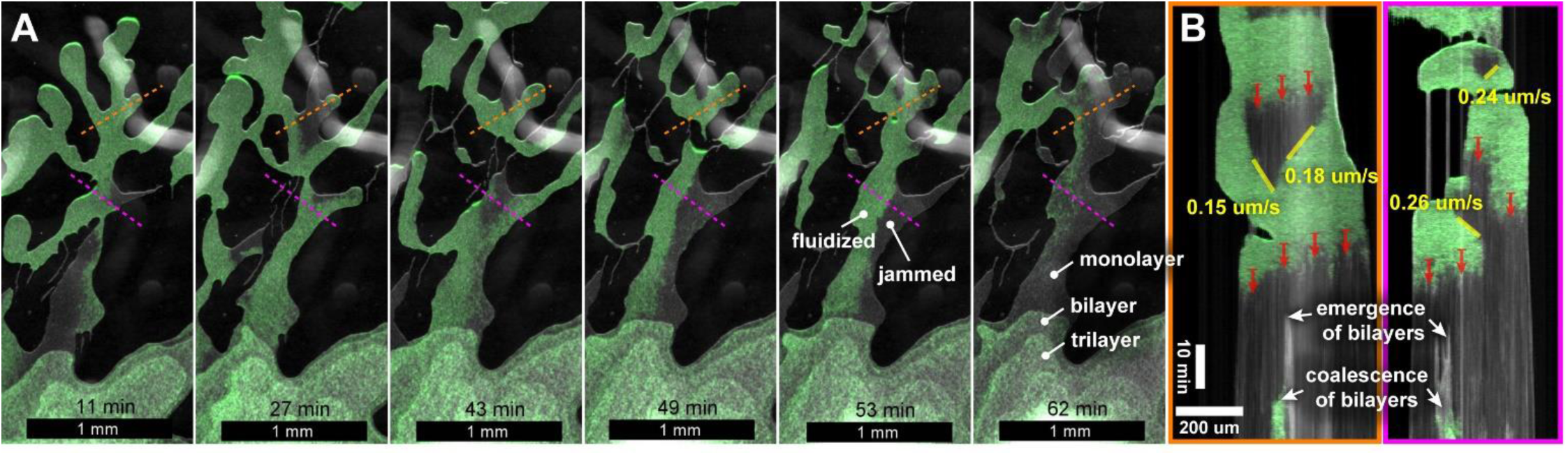
Isogenic cells exist in motile and immotile phases within the same colony. **(A)** Time-lapse imaging of wild-type *B. subtilis* spreading dendritically on a nutrient-rich soft-agarose surface. We applied a computational image filter that uses local intensity fluctuations over time to report on the degree of movement. Green regions are high motile activity and stationary regions are left gray. Transient motile regions and fluctuating boundaries between motile and stationary regions emerged and dissipated on timescales shorter than (potential) phenotypic changes (see Movie S1). **(B)** Kymographs along the colored dashed lines in (A) (time is downward). Transitions from large-scale motility (green) to immobilized (jammed) states are denoted by the red arrows. Regions could transition from jammed back to fluidized if they came in contact with a fluidized region (see boundary movements in yellow). At later times, once the entire population had become immotile, cells grew ‘upward’ into bilayers that frequently coalesced. See Movie S6.

In wild-type cells, the rapid movement of dendrites and cellular groups was correlated with significant increases in the quantitative measure of activity from our image analysis, indicating that movement of individuals positively correlates with movement of the group. Conversely, regions whose constituent cells had a low measure of activity were stationary and did not move or flow over the surface (Fig. 1 and Movie S6). Frequently, stark boundaries in activity level formed within contiguous regions of cells, indicating that within a genotypically and phenotypically identical population, sub-populations could adopt two qualitatively distinct states of motion at the same time. We interpreted the high activity state as a fluidized (low viscosity) state in which cells were swarming, and the low activity state as a ‘jammed’ state where the network of contacts and forces between cells led to local immobility. We observed that jammed regions could be fluidized if they came into contact with a fluidized region. Further, the phase boundary between these regions actively fluctuated in time, translating fractions of a micron per second (Fig. 1B), meaning that individual cells transitioned between motile and immotile states faster than (potential) phenotypic changes in the local environment. Physical theories of jamming in granular materials (61, 63, 64, 71) predict that the transition between jammed and fluidized states result from small (technically infinitesimal) differences in packing fraction.

We took two approaches to assess the connection between packing fraction and motion. Phase contrast video microscopy allowed us to visualize the cells as dark objects, while open spaces – even a fraction of a cell’s area – are relatively bright (SI Fig. 3A, Movie S7A). First, we reasoned that at suitably high magnification (60X), the average intensity in a region is directly correlated with the amount of open space (assuming the cells are in a monolayer, see next section of Results), or in other words, brighter regions have lower packing fractions. We hypothesized that if packing fraction was controlling the transition, there should be a positive correlation between local intensity and the local level of activity. We averaged both the intensity and the activity over regions ∼0.8 µm^2^, and then calculated (i) the histogram of spatially correlated values between those two measurements, (ii) the mean local intensity vs. the local activity, and (iii) the bivariate correlation coefficient (SI Fig. 3B). The mean intensity values increased monotonically with activity and the correlation coefficient was 0.34 (*p* < 10^−6^, relative to the null-hypothesis of no correlation between the two signals, SI Fig. 3B). Second, we hypothesized that there should be a negative correlation between packing fraction and flow speed – with the understanding that the nature of the jamming transition means that the relationship between those observables is non-linear (i.e. we would expect a significant, but weak negative correlation). After removing low frequency background variations, we used a fixed threshold to define which pixels were cells (dark) and which were open space (light), and averaged the resulting packing fraction over square regions 12.4 µm^2^ in area. We then used open-source particle image velocimetry software (72) to measure the local speed of cellular flow through the same pixel groups. We calculated the histogram of spatially correlated values between flow speed and packing fraction, with a bivariate correlation coefficient of – 0.11 (with *p* < 10^−6^, relative to the null-hypothesis of no correlation between the two signals) (SI Fig. 3C). Thus both methods provide significant support for the hypothesis that packing fraction regulates collective movement.

Next, we wanted to determine if available water (and its anticorrelate: packing fraction) could regulate the transition between the fluidized / collectively motile state and the jammed immotile state. We imaged expanding tips that were suddenly subjected to evaporation, and hence reduced water availability. As water evaporated, the expanding tips exhibited a dynamic transition from a fluidized and rapidly swarming state to a state where both individuals and the group were immotile (Fig. 2A/B and Movie S7B). The motile-to-immotile transition did not occur at the same time across the group, and local variations in density allowed some cells to ‘rattle’ in place, even after the group as a whole had become immotile – consistent with the appearance of ‘rattlers’ in jammed granular systems (73, 74). These observations further support the hypothesis that the transition is not a phenotypic change and that local drag and frictional forces are not preventing motion. Then, to examine the reversibility of this transition, we took the same plates and expanding tips, resealed them to halt evaporation, and continued imaging while water from the gel rehydrated the cells. Over a few minutes, the colony re-fluidized into domains of high activity sub-groups, which then coalesced until the entire tip region regained fluidity and continued to expand (Fig. 2C/D and Movie S7C). This strongly indicates that re-wetting and the corresponding reduction in packing fraction ‘reverses’ the jamming-like transition back to a fluidized and collectively motile state (61, 71, 75).

**Figure 2.**
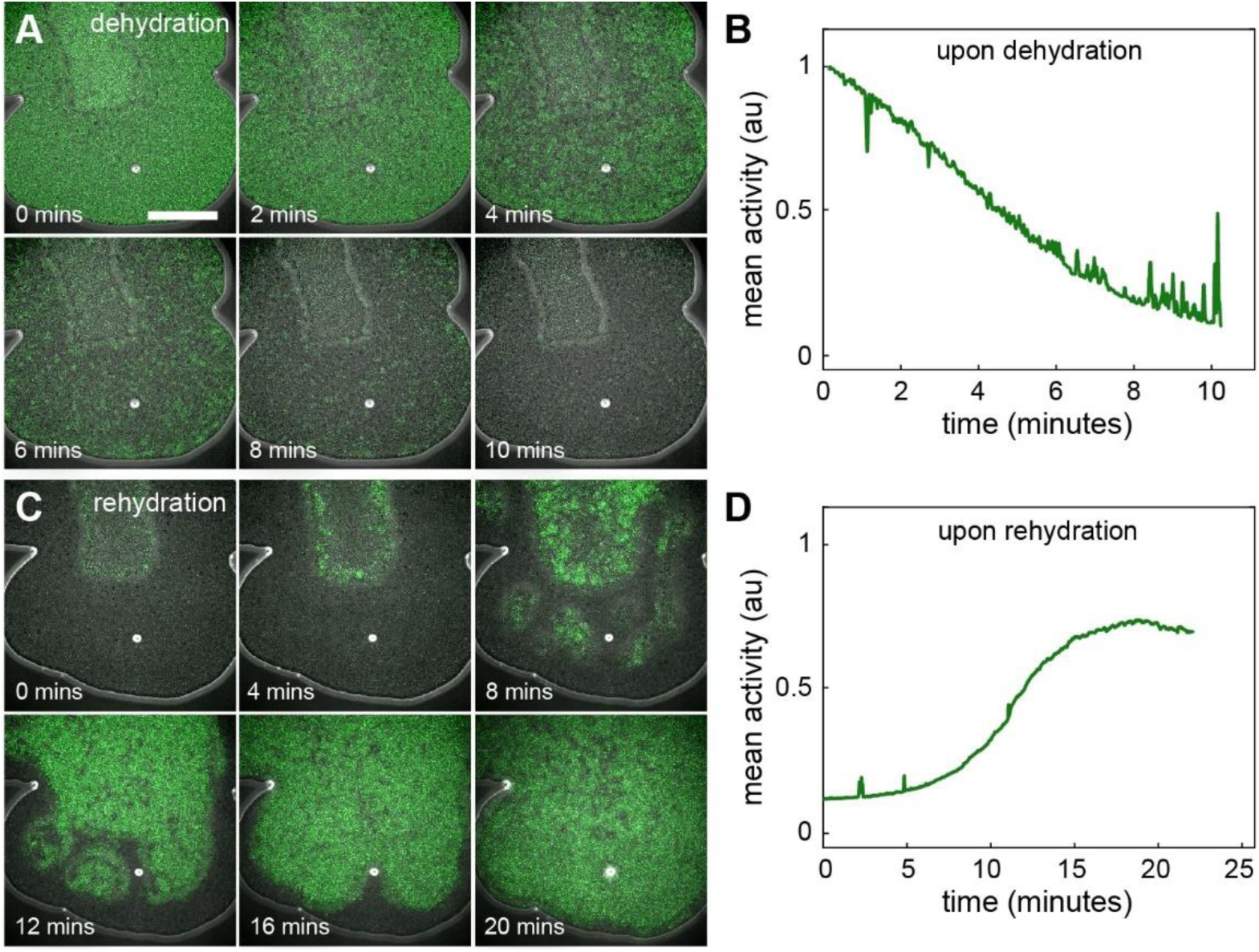
Water availability controls a jamming-like transition. **(A)** Time-lapse imaging of the tip of a wild- type *B. subtilis* dendrite on a nutrient-rich low-agarose surface. At *t* = 0 the plate was unsealed and its surface was exposed to air. Subsequent evaporation reduced water availability at the surface, causing both collective motion (green color) and the computational measure of mean motile activity **(B)** to drop as the cellular group entered the jammed state. Across the entire dendrite tip collective motility halted. **(C - D)** The same plate was then immediately resealed and imaged over 20 minutes. The punctuated drop in evaporation rate allowed cells to retain more water from the gel. Fluidized domains of motile cells subsequently emerged and coalesced until the entire dendrite was, once again, collectively motile. In both sets of images, the rectangular structure toward the top is a cellular bilayer (see Fig. 3). Noise in the activity measurement (B) was (in part) due to manual refocusing required by the retraction of the surface as water evaporated. Scale bar is 200 µm. See Movies S7A and S7B.

### Motile groups are pulled by the front in discrete layers

In our bright-field imaging it appeared that motile dendrites frequently traversed surfaces in discrete layers of cells, first traversing the surface as dense monolayers, then double layers, and so forth, up to 4 discrete layers (see Movies S6, S8 and S9). To confirm that groups migrated in discrete layers we measured the surface height of expanding dendrites with high *Z*-resolution using an interferometric profilometer (see Methods). In Fig. 3A/B we show the height field and a 1D profile for an expanding monolayer dendrite – its height across the expansion region was almost exactly 1 µm, the thickness of a single *B. subtilis* cell. As dendrites slowed and/or encountered space constraints, the layer thickness jumped to higher integer values, as shown in Fig. 3C/D whose dendrites transition between 0 and 3 layers of cells (4 layers visible in Movies S6 and S10). Looking back to the jamming data in Figs. 1 and 2, multiple discrete layers are visible in the interior of the colony. Frequently, the fastest dendrites were advancing as monolayers (the fastest moving groups in Movies S6, S8, and S9, notably tips in S10 appeared multi-layered). In addition to being valuable *in situ* data of motile-group structure, this also meant that our imaging of rapid motile groups (and attendant activity filtering) was characterizing the motion of all cells in *Z*, as opposed to there being motile and immotile cells at different *Z* positions (e.g. a biofilm blocking view of motile cells). This discrete structuring is noteworthy because relevant models of surfactant-driven spreading take cell-layer thickness as a continuous variable, whereas these data demonstrate that (frequently, though not strictly) motile groups move in monolayers with possible transitions to other discrete heights, especially when near the jamming transition.

**Figure 3.**
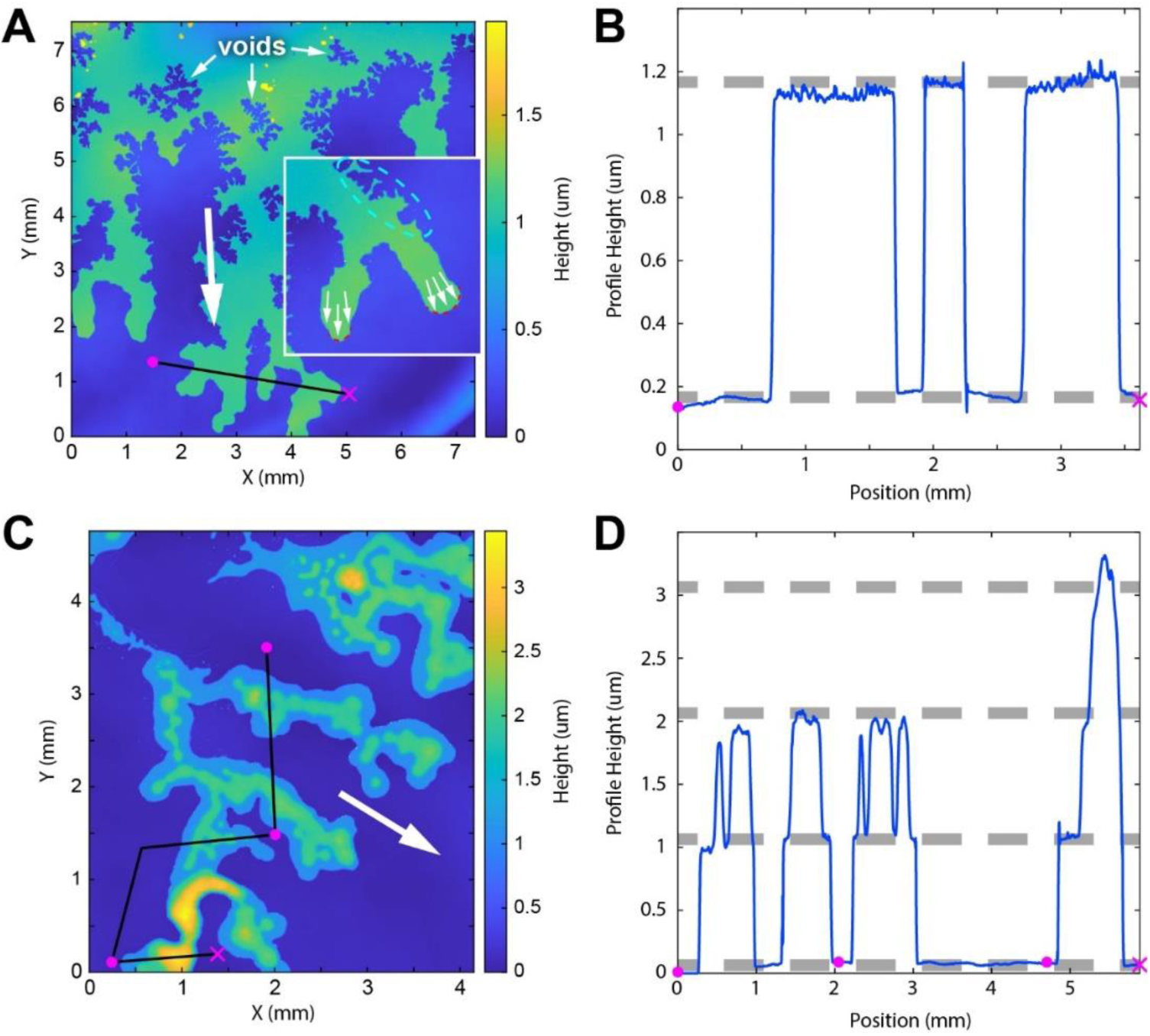
Dendrites expand in discrete layers. The left-hand column shows height images of wild-type *B. subtilis* dendrites on soft agarose taken using an interferometric profilometer (see Methods); the image backgrounds have been computationally flattened. **(A)** Advancing dendrites flowed downward (white arrow) and the fronts of those dendrites exhibited overall positive curvature (red dashed lines, inset) indicating outward pressure. Behind the advancing front, boundaries between agarose and cells exhibited a ‘lacunar’ structure (e.g. inset, cyan dashed ellipse), where the mode of the curvature distribution was negative (SI Fig. 3). Voids appeared behind the front, consistent with negative pressure in those regions overcoming meniscal forces. Both voids and lacunar structure are consistent with viscous fingering and cells de-wetting and being pulled to the front. **(B)** A one-dimensional profile of the surface height along the black line in (A). Here the advancing front is a densely packed cellular monolayer, whose height is one cell thickness as shown by the horizontal grey dashed lines (1 µm apart). **(C)** Same imaging modality as in (A), now after dendrites have jammed on the surface and then grown for ∼ 1 hr. The original flow direction is indicated by the white arrow. **(D)** A one dimensional profile of the surface height along the black line in (C). Within contiguous cellular regions, bacterial film height is discretized into layers one cell thick; here the profile shows transitions from monolayer, to bilayer, to trilayer. Discretization ceased above four layers, when presumably cells move in any orientation, rotating about their long axis within the entire vertical structure (see also Movies S6, S8 and S9).

The boundary shape between bare agarose and dense cellular monolayers also contained information. While the shape of the meniscal boundary can be influenced by surface pinning and depinning (76), this suite of imaging data showed that such boundaries have different shape statistics near and far from the advancing front. At the advancing front, the boundary curvature is smooth (low variation) with an overall positive curvature whereas behind the front retraction of material causes curvature to vary widely in a so-called ‘lacunar’ structure (SI Fig. 4A). Further, within the regions whose boundaries were lacunar, voids appeared with similar distributions of boundary curvature to the external boundary (SI Fig. 4B). Consistent with previous work on ‘viscous fingering’ (77) and models of surfactant-driven flow (23, 24), these data support the hypothesis that the advancing front has positive pressure (pushes outward, covers new territory) while the boundary behind the front experiences negative pressure (pulls inward, retracts and reduces coverage), both consistent with cellular groups primarily being front-pulled by gradients in surface tension, not pushed from the back by proliferation. Consequently, on one occasion we observed a dendrite tip contacting its own surfactant field (78), resulting in an almost immediate (within ∼ 15 s) cessation of motion (SI Fig. 5 and Movie S11) despite the number of cells in the tip increasing even after self-contact. Also consistent with a front-pull mechanism, much of the same imaging data showed independent islands of cellular monolayers, completely detached from their parent colony, traversing the surface independently, with characteristic positive curvature at the front and negative curvature regions in the back (Movies S6, S8, S12 and S13).

While high numbers of cells are required for group motility, these data indicate that proliferation alone is not generating outward pressure to drive group motion. To definitely determine this, we measured the area of dense monolayer islands of cells as they moved over time. In Fig. 4, we show one such island traversing an agarose surface over a distance many times its length. Rather than increasing due to cell division and growth, the island area decreased over time as such islands tend to leave a trail of cells immobilized on the surface by the meniscus. Therefore, during surfactin-driven surface motility groups of cells are capable of large-scale motion without growth in the population and without connection to their parent colony. Rather, boundary curvature analysis and the existence of independent islands provide evidence that cells are pulled by the front, frequently in monolayers at a density near the threshold for jamming.

**Figure 4.**
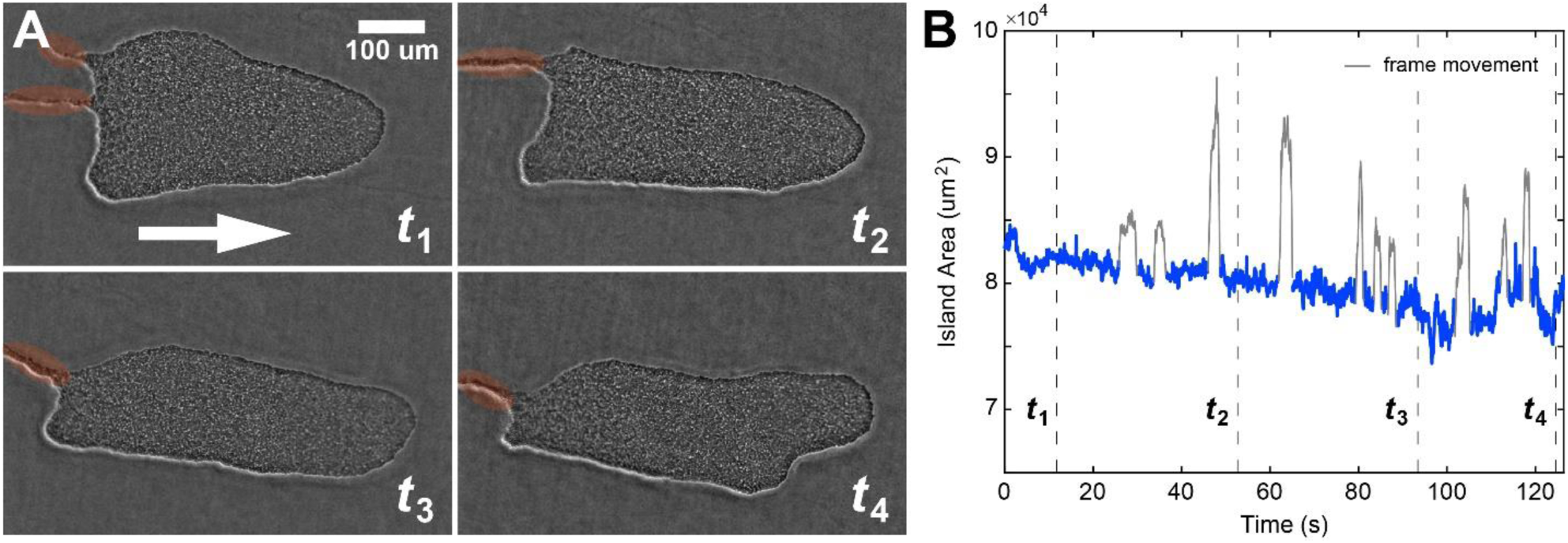
Groups can move independently without growth. **(A)** Four snapshots of an independent monolayer ‘island’ of wild-type *B. subtilis* cells moving to the right (white arrow) on soft agarose. Over the course of imaging, the group moved at an average speed of ∼ 6 µm/s and the total amount of movement was ∼ 700 µm, necessitating movement of the viewing frame (grey data in B). Common to our observations, such islands leave a trail of cells immobilized on the surface behind them (orange highlighted regions). **(B)** The area of the island decreased over time (time labels correspond between A and B). When combined with our data showing movement in monolayers, these data demonstrate that movement does not require proliferation. See Movie S13.

### Material flows from parent colony to dendrite tip

While our data were consistent with outward front-pressure pulling the group away from the parent colony, multiple time-lapse experiments showed apparent rapid flow of discrete cellular layers from parent colony to the expanding dendrite tip (e.g. see Movie S10). To confirm that these observations were indeed flow of material over the scale of millimeters to centimeters, we performed our typical wild-type expansion assay, but doped the initial cellular deposition with 1 µm fluorescent polystyrene beads, at a ratio of ∼ 1 bead per 100 cells (see Methods). We imaged the motion of the tracer beads at 30 fps, which allowed us to see the trajectories of individual beads on the time scale of minutes (Movie S14, SI Fig. 6A). The swarming motility of wild-type cells caused the motion of individual beads to be erratic, akin to a non-thermal random walk with drift (79) especially when flow speed was slow in comparison to cellular swim speeds (Movie S15, SI Fig. 6B). This erratic motion prohibited flow tracking in wild-type groups, however, as we had confirmed earlier (38), mutants of *B. subtilis* unable to perform flagellar self-propulsion (Δ*hag*) exhibit rapid surface motility with speeds and morphologies similar to wild-type cells (see SI Fig. 2). This strain did not cause severe bead agitation and thus we were able to perform *in situ* flow measurements using those mutants (Fig. 5 and Movie S16).

**Figure 5.**
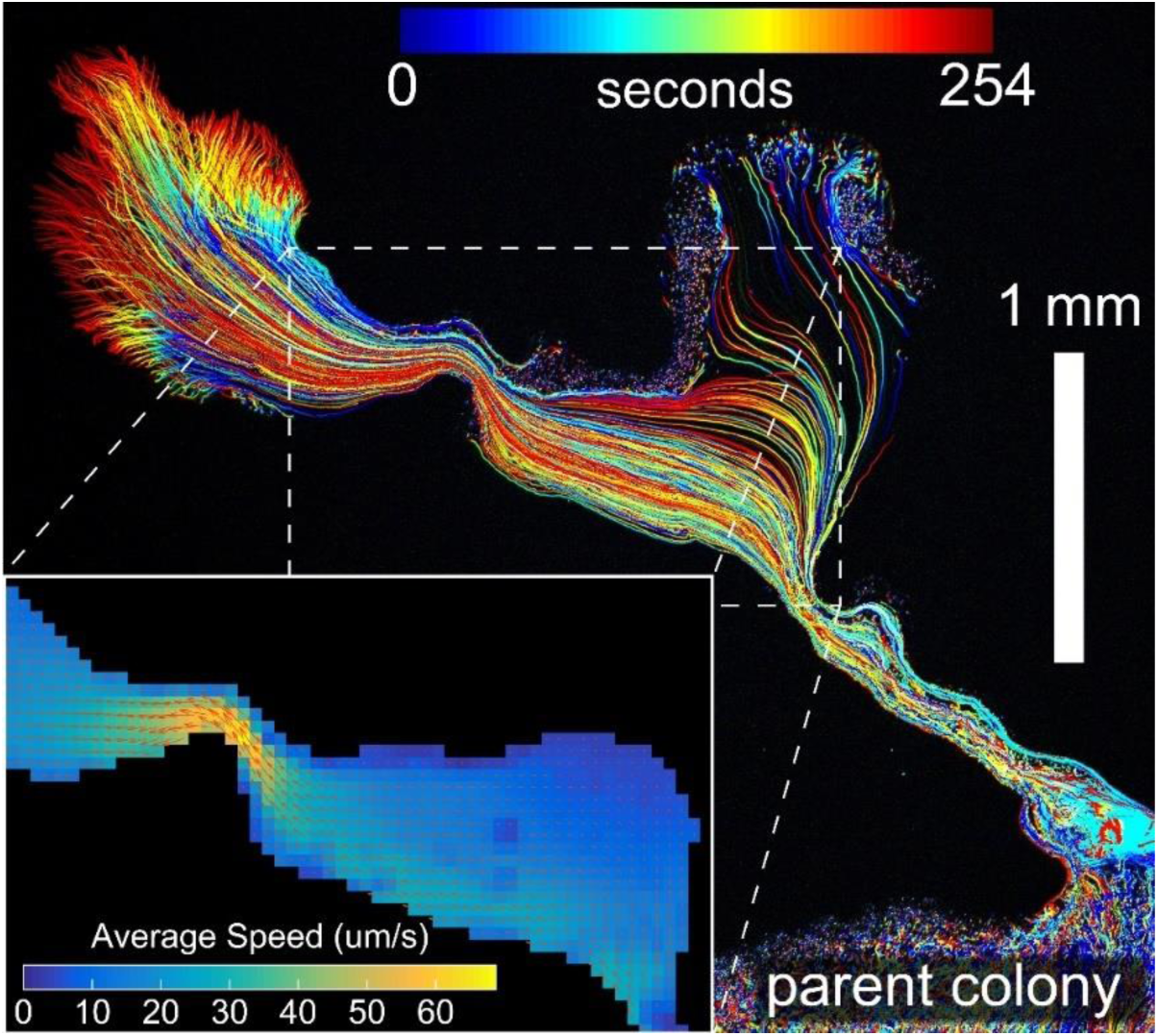
Dendrite expansion corresponds with large-scale flow. Fluorescent 1 µm tracer beads were diluted with the cellular suspension and deposited onto the agarose surface. We examined the motion of beads in Δ*hag* mutant cells (lacking flagella); this minimized bead agitation due to individual cellular motility (see SI Fig. 5 and Movies S14 and 15 for wild-type). The beads were imaged in an expanding dendrite millimeters long on the time scale of minutes; the positions of individual beads are shown colored by time. Here the dendrite is connected to its parent colony, material flow moved beads from the parent colony into the dendrite. The accompanying movie (Movie S16) shows individual beads starting in the colony and moving to the dendrite tip, demonstrating that flow generated by pulling at the front can move material long distances compared to cell size, from the colony to the tip of a growing dendrite. The inset shows the average flow velocity (orange arrows) and average flow speed through a region with varying width. Similar to a 2D incompressible fluid, constrictions accelerated flow, whereas flow slowed at wider points along the dendrite.

Using open-source cross-correlation software, we measured bead flow via particle image velocimetry (72). Similar to the flow characteristics of a typical incompressible fluid, the flow speed of dense cellular monolayers decreased in regions with wide meniscal boundaries and significantly sped up at points of constriction (see inset Fig. 5). The similarity between incompressible 2D flow and flow of dense cellular monolayers is consistent with the fact that cells are effectively solid steric objects densely packed in (incompressible) water. Average flow speeds, and following the motion of individual beads, showed that material originating in the parent colony frequently made its way to the expanding tip (Fig. 5 and Movie S16). Thus, while our island data showed that motile groups need not be connected to the parent colony and that cellular proliferation was not required for group motility, these data indicate that when connected back to the parent colony, cells and material (e.g. nutrients) can be supplied to the expanding tip of a dendrite.

Having confirmed that our bright-field data was indeed revealing large scale flow, we examined how the speed of tip expansion into fresh territory compared with the speed of cellular flow along established dendrites connecting the tip to the parent colony (SI Fig. 7). Consistent with the bead-flow data, kymograph analysis showed that when expanding dendrites are connected to their parent colony, cells (and surrounding fluid) flow many centimeters from the colony into the expansion zone. Such flows along an established dendrite are significantly faster (∼ 10 µm/s) than the speed of tip movement (∼ 3 µm/s) into fresh territory, as they must be for material to reach the tip from the colony. At this point it is unclear how groups create these differentials in speed along a dendrite, but we offer a potential mechanism in the Discussion.

### Motile groups use material flow to traverse nutrient poor regions

We knew from previous work (37, 38) that neither chemotaxis nor flagellar-mediated motility is required for group surface motility, but that surface motility was potently modulated by (e.g.) the presence of monovalent ions (K+ (25)). This suggested that groups might respond to their chemical environment indirectly via changes in metabolism and/or gene expression. For instance, differences in nutrient concentration might affect the rate of synthesis and/or secretion of surfactin, and thereby affect surfactin-mediated motility. Further, our large-scale flow data suggested that differences across the chemical landscape that affect group surface motility could be ameliorated by flow of material from other regions.

We wanted to test whether motile groups had differential responses to exogenously presented nutrient gradients, and specifically whether groups would avoid negative nutrient gradients. We created agarose plates with two distinct halves – one half containing the same rich defined medium (RDM) used earlier, the other half containing (potassium-free) NaCl buffer osmotically matched with the RDM to maintain the same osmotic potential (see Methods). A thin impermeable barrier separated the two halves, initially keeping the nutrients on one side. Once the agarose set, we poured a thin (∼ 1 mm) layer of osmotically matched NaCl buffer with agarose, and thus linked the two regions into a contiguous surface with uniform mechanical properties that allowed cells to move freely between them. The nutrient-rich half also contained a red fluorescent tracer dye (rhodamine) that served as an approximate reporter of nutrient diffusion over the barrier. With or without cells, these plates showed diffusion of the dye from the nutrient rich side into the side devoid of nutrients, meaning that there was a negative concentration gradient on the surface from the nutrient rich to the nutrient poor side. We reasoned that if wild-type cells were responding (potentially indirectly) to the outward-facing chemical gradients on a standard plate (i.e. with initially isotropic nutrients) or to the absence of required potassium ions on the split-plate, they would react to the reverse nutrient gradient on these split plates and either avoid the nutrient poor region or show reduced motility toward it.

To test this, we simultaneously inoculated such a split-plate in two positions from the same isogenic population of wild-type cells, one colony on the nutrient rich side, one on the nutrient poor side (Fig. 6A/B). The colony inoculated on the nutrient poor side did not exhibit collective motility and was effectively stationary, while the colony on the nutrient rich side rapidly covered the nutrient rich zone and simultaneously ventured into the nutrient poor zone. Initially, dendrites that expanded into the nutrient poor zone covered the plate less densely than dendrites that never left the nutrient rich zone (Fig. 6B), but tip speeds were similar in both zones. Ultimately, cells originating from the nutrient-rich zone colonized the entire plate (both zones, see Movie S17). The difference in initial coverage between the zones supports the hypothesis that groups do exhibit a (mild) chemotropic response to gradients in nutrients, ions, or potentially their own secreted waste products. Interestingly, these data show that connection to a parent colony in a nutrient rich zone allows cells to explore nutrient-poor regions, even if those regions lack critical chemical species (K+ ions in wild-type (25)). This also complements findings that bacteria can transport other bacteria as cargo during surface motility (80). We did not observe any independently moving islands of cells in the nutrient-poor region. Our flow data offer a qualitative mechanism for this ‘scouting’ ability, that large-scale flow of material from parent colony to the extending tip brings metabolic resources, presumably to fuel, in part, surfactin production, thus allowing Marangoni forces to continue to pull the group outward via positive tip pressure.

**Figure 6.**
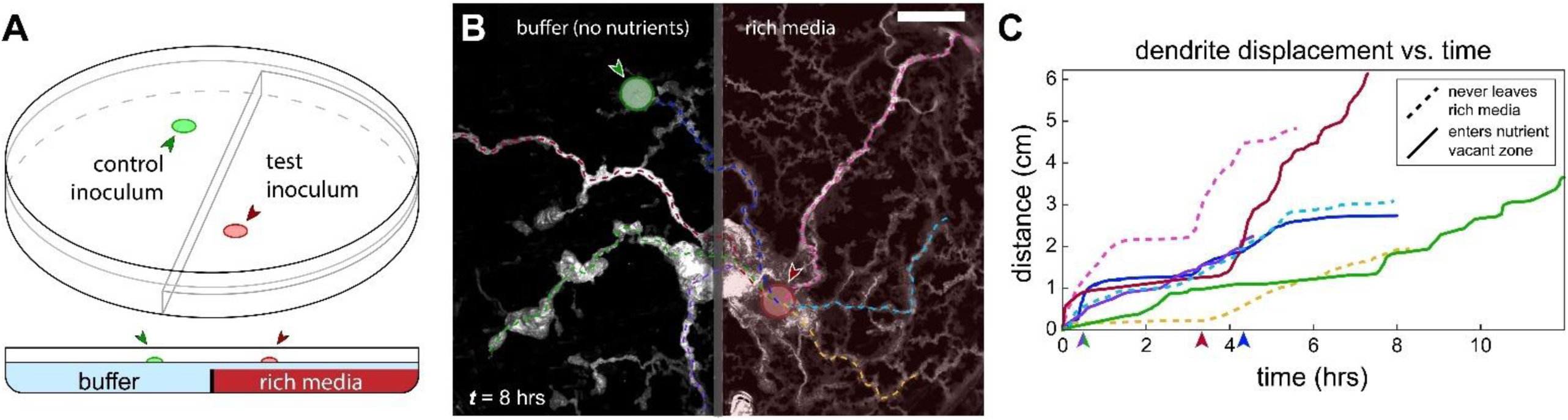
Surface motility in the presence of an exogenous nutrient gradient. **(A)** Soft agarose plates were poured with a barrier between nutrient-rich media doped with a fluorescent tracer dye (rhodamine) on the right half of the plate and osmotically balanced zero-nutrient buffer on the left, with a top layer of balanced zero-nutrient buffer to create a contiguous surface. The spatial distribution of the dye confirmed nutrient diffusion with a negative nutrient gradient from right to left. Wild-type *B. subtilis* colonies were deposited on either side of the barrier. **(B)** The colony inoculated in the nutrient-rich zone (right, red circle) spread dendritically across the divided plate into the zero-nutrient zone, effectively traversing along a negative nutrient gradient. An identical inoculum in the zero-nutrient zone (left, green circle) exhibited no collective motility. Scale bar is 1 cm. **(C)** Some dendrites remained in the nutrient-rich zone (dashed lines) while other dendrites ventured into the zero-nutrient zone (solid lines). Both classes of dendrites exhibited similar speeds, and flow moved material along dendrites in both regions (see also Movie S17). The small arrows on the time axis are when each dendrite entered the zero-nutrient zone. Colors are matched between (B) and (C).

## Discussion

Our data support a model in which water availability – a proxy for bacterial packing fraction – is a sensitive control parameter for an abiotic jamming transition in the granular material that is densely packed, expanding bacterial populations. We hypothesize that motility plays a contributory role, encouraging a fluidized state via motility-dependent viscosity (59, 81). When sufficient fluid is extracted from the substrate by endogenous osmolyte secretion and cells are in a fluidized state, groups flow over the surface driven and guided by surface-tension gradients (Marangoni forces). Conversely, when water availability decreases, either by (e.g.) evaporation or decreasing osmotic potential, packing fraction increases and cells transition to a jammed and immotile state. It is worth noting that while our data are consistent with a jamming-fluidization transition, we cannot use our current data to *prove* that we are meeting the technical requirements of such a transition. Jamming is defined by a non-crystalline network of steric contacts and forces between particles that are ‘isostatic’, that is, they do not have free modes of motion (other than rattlers (73)), but we are unaware of a modality of microscopy that permits live, *in situ* visualization of the force and contact network in a dense bacterial group (stationary or collectively motile).

Nonetheless, the potential existence of an abiotic jamming transition that regulates group motility through the dense packing of nematic actors offers a mechanism for why small increases in agarose gel stiffness (∼0.05%) switch groups from rapid surface migration to slow steric growth. It also clarifies why the secretion of both surfactants and osmolytes is required for collective motility (37, 50); the former being crucial for force generation and the latter for maintaining fluidization. Thus we hypothesize that colonies that are not able to achieve the fluidized state – for any reason – are constrained to expand on the order of 100 times slower over surfaces using the forces generated by cell wall growth and cell division (i.e. the mode of sessile growth one typically associates with single colonies growing on a petri dish).

Our examination of boundary structure (SI Fig. 4) was consistent with outward-facing pressure at the expanding front and inward-facing pressure behind the front. However, many data sets showed persistent flow from parent colony, or other regions behind the front, toward the advancing front, without the appearance of lacunar structure (e.g. Movies S2 and S10). We speculate that these differences in boundary structure, and correspondingly flow, might reflect differences in the magnitude of force being generated at the front and/or changes to the effective viscosity of the cellular suspension. Differences in force magnitude might arise from varying rates of surfactin secretion and/or processes that affect the spreading or degradation of surfactin. Differences in effective viscosity of the cellular suspension could arise due to changes in cellular density (including but not limited to jamming), cellular aspect ratio (9, 82), internal shear arising from individual motility (59, 81), or processes that affect the rate of fluid extraction (e.g. via osmolytes) or loss (e.g. via evaporation) from the substrate. Similarly, we hypothesize that the difference in tip speed vs. flow speed along an established dendrite (SI Fig. 7) – which allows material to be brought from parent colony to dendrite tip – might reflect different energetic costs for motion in those two scenarios. A tip advancing into fresh territory uses surface tension differences to generate force, but, given our layering data, it must also pay a line-energy cost along the meniscus on each side of the dendrite, which to first order would appear like a constant cost in energy per unit length of movement (equivalent to a constant reduction in overall force). Conversely, assuming that cells flowing within an established dendrite are using the same mechanisms for force generation, they move within the same meniscal boundaries established by the tip, thus do not pay this energetic cost, and thus have a larger effective force pulling them outward.

The existence of large scale flow from parent colony to dendrite tip presents a number of biophysical questions and possibilities. How far into nutrient poor zones can expanding dendrites venture? How do those distances depend on the available nutrients or the position of the parent colony? How do surface properties like local water availability and topography (83, 84) affect flow and colonization? How do structures on the surface guide and/or impeded collective motility? The fact that we observed islands of cells moving independently on nutrient-rich surfaces, whereas flow from a parent colony was required for exploration of nutrient-poor zones, suggests that those modalities of motion (island vs. flowing dendrite) depend on environmental conditions. For instance, an island that ventures into a nutrient poor region might quickly deplete its resources, and thus decrease secretion of osmolytes and surfactants required for motion (54). Likewise, the transport of materials and nutrients from a common pool (parent colony) into discrete tendrils far from the common pool (80) spur game-theoretic questions of how those resources are divided, whether phenotypic changes are playing a role, and – like other collectively motile and reproductive systems (e.g. *Dictyostelium* (85, 86)) – how the specific genomes of individuals propagate in space and time relative to their nearly isogenic kin.

Whereas individual cells in bulk fluids propel themselves and direct their motion according to the algorithms of chemotaxis, these data and previous work strongly suggest a fundamentally different set of rules govern group surface motility. Specifically, that collective secretion of surfactants and osmolytes drives and directs motion, that large-scale flow plays a critical role in exploration of territories with differing nutrient concentrations, and that bacterial packing fraction – modulated through multiple mechanisms – is a sensitive control parameter that regulates transitions between fluidized (collectively motile) and jammed (immotile) states of the group. Thus continued understanding and modeling of microbial communities on surfaces must consider how these physical factors – not fully described by genetics and biochemistry – contribute to population dynamics.

## Methods

### B. subtilis culture storage and preparation

Isogenic cultures of *B. subtilis* strains were grown to OD 0.5, mixed 50% in glycerol aliquots, individually snap frozen and stored at - 80 C. For each experiment, 200 µL aliquots were removed and thawed, then diluted in 10 ml LB medium and grown in a shaking incubator at 37 C for 4 hours until mid-log phase, then pelleted by centrifugation at 4000 g for 10 mins. The supernatant was removed, and the bacterial pellets were then resuspended in 200 µL LB, yielding a culture with an approximate density OD 10, then 1.5 µL of that culture was deposited on 0.5% agarose plates with Teknova EZ-RDM media (a rich defined medium), then dried for 10 mins in air and sealed with parafilm. For bead flow experiments, 1 µm fluorescent polystyrene beads (Bangs Laboratories, Dragon Green) were diluted into the inoculation culture and vortexed before deposition.

Depending on the type of time lapse experiment, plates were then either incubated for 1 hr then imaged or imaged immediately at 37 C. For culture profilometry (height measurements), cultures were incubated for 2 - 3 hrs at 37 C, then unsealed within 10 minutes of imaging to minimize evaporation and jamming. For the multilayer height measurements (Fig. 3B), cultures were unsealed, transitioned to immotility, and were then imaged between 60 to 80 minutes later.

### Plate preparation

Plates were created by mixing 100 mL Teknova EZ-RDM and 0.5% agarose by weight and autoclaving for 15 minutes at 121 C. Media was cooled to 50 C in an incubator before being poured into 4 plates (25 ml each). Each plate was cooled and solidified under flame in open air for 10, 15, or 30 minutes before immediate inoculation, creating a range of initial gel hydrations to examine the effects of water availability on colony morphology. For the split-plate experiment (Fig. 6) zero-nutrient buffer was made from de-ionized water, 0.5% agarose w/v and NaCl concentration matched to the calculated osmotic potential of the rich defined medium.

### Microscopy and Imaging

Transmitted-light and oblique-illumination microscopy were performed using a Nikon SMZ-25 stereo zoom microscope, with a P2-SHR Plan Apo 1x objective, and a Prior ES111 OptiScan stage. High speed images were taken using an Andor Zyla 5.5 CMOS camera. Phase contrast microscopy was performed using a Nikon Eclipse Ti-E inverted microscope with an CFI S Plan Fluor ELWD 20x Ph1 ADM objective (Fig. 2) or CFI S Plan Fluor ELWD 40X Ph2 ADM objective (SI Fig. 3). High speed images were taken using an Andor iXon Ultra 888 EMCCD camera. Culture height images were captured using a Zygo NewView 7300 optical 3D profilometer in the University of Oregon’s CAMCOR facility. Low-frequency variations in plate height were computationally removed by subtraction of a fitted a polynomial surface to the background.

Activity overlays were generated using custom Matlab© scripts which calculated the per-pixel sum over the absolute differences between 5 or 7 consecutive frames. Kymographs were generated using custom Matlab scripts to identify and interpolate contours, then sum over cross-sectional slices of intensities along the contours over time. All scripts and Matlab code are available on request.

## Acknowledgements

We thank Prof. Daniel Kearns and his lab for their generous gift of the *B. subtilis* strains used in this study and Eric Corwin for critical reading of the manuscript. This work was supported by the University of Oregon.

## Notes

#### Summary of Updates

Substantial reworking. More, better data supporting jamming transition.

## References

1. R. Mesibov, J. Adler, Chemotaxis Toward Amino Acids in Escherichia coli. J. Bacteriol. 112, 315–326 (1972).

2. L. F. Garrity, G. W. Ordal, Chemotaxis in Bacillus subtilis: how bacteria monitor environmental signals. Pharmacol. Ther. 68, 87–104 (1995).

3. D. S. Bischoff, G. W. Ordal, Bacillus subtilis chemotaxis: a deviation from the Escherichia coli paradigm. Mol. Microbiol. 6, 23–28 (1992).

4. H. C. Berg, D. A. Brown, Chemotaxis in Escherichia coli analysed by Three-dimensional Tracking. Nature 239, 500 (1972).

5. C. V. Rao, G. D. Glekas, G. W. Ordal, The three adaptation systems of Bacillus subtilis chemotaxis. Trends Microbiol. 16, 480–487 (2008).

6. R. Balagam, O. A. Igoshin, Mechanism for Collective Cell Alignment in Myxococcus xanthus Bacteria. PLOS Comput. Biol. 11, e1004474 (2015).

7. D. B. Kearns, L. J. Shimkets, Chemotaxis in a gliding bacterium. Proc. Natl. Acad. Sci. 95, 11957–11962 (1998).

8. A. E. Patteson, P. E. Arratia, A. Gopinath, Quenching a swarm: Effect of light exposure on suppression of collective motility in swarming Serratia marcescens. bioRxiv, 331801 (2018).

9. A. Rabani, G. Ariel, A. Be’er, Collective Motion of Spherical Bacteria. PLOS ONE 8, e83760 (2013).

10. T. Matsuyama, et al., Dynamic Aspects of the Structured Cell Population in a Swarming Colony of Proteus mirabilis. J. Bacteriol. 182, 385–393 (2000).

11. H. H. Tuson, M. F. Copeland, S. Carey, R. Sacotte, D. B. Weibel, Flagellum Density Regulates Proteus mirabilis Swarmer Cell Motility in Viscous Environments. J. Bacteriol. 195, 368–377 (2013).

12. B. V. Jones, Role of swarming in the formation of crystalline Proteus mirabilis biofilms on urinary catheters. J. Med. Microbiol. 54, 807–813 (2005).

13. E. Ben-Jacob, I. Cohen, H. Levine, Cooperative self-organization of microorganisms. Adv. Phys. 49, 395–554 (2000).

14. E. Ben-Jacob, From snowflake formation to growth of bacterial colonies II: Cooperative formation of complex colonial patterns. Contemp. Phys. 38, 205–241 (1997).

15. S. Atis, B. T. Weinstein, A. W. Murray, D. R. Nelson, Microbial Range Expansions on Liquid Substrates. Phys. Rev. X 9, 021058 (2019).

16. T. Ursell, R. M. W. Chau, S. Wisen, D. Bhaya, K. C. Huang, Motility Enhancement through Surface Modification Is Sufficient for Cyanobacterial Community Organization during Phototaxis. PLoS Comput. Biol. 9 (2013).

17. N. Schuergers, et al., Cyanobacteria use micro-optics to sense light direction. eLife 5, e12620 (2016).

18. M. Driscoll, et al., Unstable fronts and motile structures formed by microrollers. Nat. Phys. 13, 375–379 (2017).

19. M. F. Copeland, D. B. Weibel, Bacterial Swarming: A Model System for Studying Dynamic Self-assembly. Soft Matter 5, 1174–1187 (2009).

20. R. M. Harshey, Bacterial Motility on a Surface: Many Ways to a Common Goal. Annu. Rev. Microbiol. 57, 249–273 (2003).

21. A. Be’er, G. Ariel, A statistical physics view of swarming bacteria. Mov. Ecol. 7, 9 (2019).

22. C. Giverso, M. Verani, P. Ciarletta, Branching instability in expanding bacterial colonies. J. R. Soc. Interface 12, 20141290 (2015).

23. S. Trinschek, K. John, U. Thiele, Modelling of surfactant-driven front instabilities in spreading bacterial colonies. Soft Matter 14, 4464–4476 (2018).

24. S. Srinivasan, C. N. Kaplan, L. Mahadevan, A multiphase theory for spreading microbial swarms and films. eLife 8, e42697 (2019).

25. R. F. Kinsinger, M. C. Shirk, R. Fall, Rapid Surface Motility in Bacillus subtilis Is Dependent on Extracellular Surfactin and Potassium Ion. J. Bacteriol. 185, 5627–5631 (2003).

26. D. Kaiser, Bacterial Swarming: A Re-examination of Cell-Movement Patterns. Curr. Biol. 17, R561–R570 (2007).

27. J. D. Partridge, R. M. Harshey, Swarming: Flexible Roaming Plans. J. Bacteriol. 195, 909–918 (2013).

28. T. A. Witten, L. M. Sander, Diffusion-Limited Aggregation, a Kinetic Critical Phenomenon. Phys. Rev. Lett. 47, 1400–1403 (1981).

29. J. Y. Wakano, S. Maenosono, A. Komoto, N. Eiha, Y. Yamaguchi, Self-Organized Pattern Formation of a Bacteria Colony Modeled by a Reaction Diffusion System and Nucleation Theory. Phys. Rev. Lett. 90, 258102 (2003).

30. I. Golding, Y. Kozlovsky, I. Cohen, E. Ben-Jacob, Studies of bacterial branching growth using reaction– diffusion models for colonial development. Phys. Stat. Mech. Its Appl. 260, 510–554 (1998).

31. E. Tamar, M. Koler, A. Vaknin, The role of motility and chemotaxis in the bacterial colonization of protected surfaces. Sci. Rep. 6, 19616 (2016).

32. N. C. Caiazza, R. M. Q. Shanks, G. A. O’Toole, Rhamnolipids Modulate Swarming Motility Patterns of Pseudomonas aeruginosa. J. Bacteriol. 187, 7351–7361 (2005).

33. C. J. Ingham, E. B. Jacob, Swarming and complex pattern formation in Paenibacillus vortex studied by imaging and tracking cells. BMC Microbiol. 8, 36 (2008).

34. J. K. Parrish, L. Edelstein-Keshet, Complexity, Pattern, and Evolutionary Trade-Offs in Animal Aggregation. Science 284, 99–101 (1999).

35. E. B. Steager, C.-B. Kim, M. J. Kim, Dynamics of pattern formation in bacterial swarms. Phys. Fluids 20, 073601 (2008).

36. A. Marrocco, et al., Models of Self-Organizing Bacterial Communities and Comparisons with Experimental Observations. Math. Model. Nat. Phenom. 5, 148–162 (2010).

37. D. B. Kearns, R. Losick, Swarming motility in undomesticated Bacillus subtilis. Mol. Microbiol. 49, 581–590 (2003).

38. R. Fall, D. B. Kearns, T. Nguyen, A defined medium to investigate sliding motility in a Bacillus subtilis flagella-less mutant. BMC Microbiol. 6, 31 (2006).

39. S. Mukherjee, D. B. Kearns, The structure and regulation of flagella in Bacillus subtilis. Annu. Rev. Genet. 48, 319–340 (2014).

40. R. A. Calvo, D. B. Kearns, FlgM Is Secreted by the Flagellar Export Apparatus in Bacillus subtilis. J. Bacteriol. 197, 81–91 (2015).

41. T. S. Murray, B. I. Kazmierczak, Pseudomonas aeruginosa Exhibits Sliding Motility in the Absence of Type IV Pili and Flagella. J. Bacteriol. 190, 2700–2708 (2008).

42. A. Toguchi, M. Siano, M. Burkart, R. M. Harshey, Genetics of Swarming Motility in Salmonella enterica Serovar Typhimurium: Critical Role for Lipopolysaccharide. J. Bacteriol. 182, 6308–6321 (2000).

43. M. Burkart, A. Toguchi, R. M. Harshey, The chemotaxis system, but not chemotaxis, is essential for swarming motility in Escherichia coli. Proc. Natl. Acad. Sci. U. S. A. 95, 2568–2573 (1998).

44. H. Du, et al., High Density Waves of the Bacterium Pseudomonas aeruginosa in Propagating Swarms Result in Efficient Colonization of Surfaces. Biophys. J. 103, 601–609 (2012).

45. A. Yang, W. S. Tang, T. Si, J. X. Tang, Influence of Physical Effects on the Swarming Motility of Pseudomonas aeruginosa. Biophys. J. 112, 1462–1471 (2017).

46. R. Chen, S. B. Guttenplan, K. M. Blair, D. B. Kearns, Role of the sD-Dependent Autolysins in Bacillus subtilis Population Heterogeneity. J. Bacteriol. 191, 5775–5784 (2009).

47. A. Be’er, et al., Paenibacillus dendritiformis Bacterial Colony Growth Depends on Surfactant but Not on Bacterial Motion. J. Bacteriol. 191, 5758–5764 (2009).

48. E. Ward, et al., Organization of the flagellar switch complex of Bacillus subtilis. J. Bacteriol., JB.00626-18 (2018).

49. J. R. Kirby, T. B. Niewold, S. Maloy, G. W. Ordal, CheB is required for behavioural responses to negative stimuli during chemotaxis in Bacillus subtilis. Mol. Microbiol. 35, 44–57 (2000).

50. D. B. Kearns, A field guide to bacterial swarming motility. Nat. Rev. Microbiol. 8, 634–644 (2010).

51. L. W. Schwartz, R. V. Roy, Some results concerning the potential energy of interfaces with nonuniformly distributed surfactant. Phys. Fluids 13, 3089–3092 (2001).

52. Á. T. Kovács, Bacillus subtilis. Trends Microbiol. 27, 724–725 (2019).

53. T. E. Angelini, M. Roper, R. Kolter, D. A. Weitz, M. P. Brenner, <em>Bacillus subtilis</em> spreads by surfing on waves of surfactant. Proc. Natl. Acad. Sci. 106, 18109 (2009).

54. W.-J. Ke, Y.-H. Hsueh, Y.-C. Cheng, C.-C. Wu, S.-T. Liu, Water surface tension modulates the swarming mechanics of Bacillus subtilis. Front. Microbiol. 6 (2015).

55. S. Srinivasan, N. C. Kaplan, L. Mahadevan, Dynamics of spreading microbial swarms and films. bioRxiv (2018) https:/doi.org/10.1101/344267 (February 5, 2019).

56. M. Fauvart, et al., Surface tension gradient control of bacterial swarming in colonies of Pseudomonas aeruginosa. Soft Matter 8, 70–76 (2012).

57. H. H. Wensink, et al., Meso-scale turbulence in living fluids. Proc. Natl. Acad. Sci. 109, 14308–14313 (2012).

58. H. M. López, J. Gachelin, C. Douarche, H. Auradou, E. Clément, Turning Bacteria Suspensions into Superfluids. Phys. Rev. Lett. 115, 028301 (2015).

59. A. Sokolov, I. S. Aranson, Reduction of Viscosity in Suspension of Swimming Bacteria. Phys. Rev. Lett. 103, 148101 (2009).

60. M. Burel, S. Martin, O. Bonnefoy, Jamming/flowing transition of non-Brownian particles suspended in a iso-density fluid flowing in a 2D rectangular duct. EPJ Web Conf. 140, 03086 (2017).

61. E. Woldhuis, V. Chikkadi, M. S. van Deen, P. Schall, M. van Hecke, Fluctuations in flows near jamming. Soft Matter 11, 7024–7031 (2015).

62. K. A. Dahmen, Y. Ben-Zion, J. T. Uhl, A simple analytic theory for the statistics of avalanches in sheared granular materials. Nat. Phys. 7, 554–557 (2011).

63. M. E. Cates, J. P. Wittmer, J.-P. Bouchaud, P. Claudin, Jamming, Force Chains, and Fragile Matter. Phys. Rev. Lett. 81, 1841–1844 (1998).

64. A. J. Liu, S. R. Nagel, Jamming is not just cool any more. Nature 396, 21–22 (1998).

65. I. R. Peters, et al., Dynamic jamming of iceberg-choked fjords. Geophys. Res. Lett. 42, 1122–1129 (2015).

66. J. Henrichsen, Bacterial surface translocation: a survey and a classification. Bacteriol. Rev. 36, 478–503 (1972).

67. N. Morales-Soto, et al., Preparation, Imaging, and Quantification of Bacterial Surface Motility Assays. J. Vis. Exp. JoVE (2015) https:/doi.org/10.3791/52338 (October 7, 2019).

68. A. J. Wolfe, H. C. Berg, Migration of bacteria in semisolid agar. Proc. Natl. Acad. Sci. 86, 6973–6977 (1989).

69. L. Jauffred, R. Munk Vejborg, K. S. Korolev, S. Brown, L. B. Oddershede, Chirality in microbial biofilms is mediated by close interactions between the cell surface and the substratum. ISME J. 11, 1688–1701 (2017).

70. C. Beloin, A. Roux, J.-M. Ghigo, Escherichia coli biofilms. Curr. Top. Microbiol. Immunol. 322, 249–289 (2008).

71. C. Reichhardt, C. J. O. Reichhardt, Aspects of jamming in two-dimensional athermal frictionless systems. Soft Matter 10, 2932–2944 (2014).

72. W. Thielicke, E. Stamhuis, PIVlab – Towards User-friendly, Affordable and Accurate Digital Particle Image Velocimetry in MATLAB. J. Open Res. Softw. 2, e30 (2014).

73. M. Skoge, A. Donev, F. H. Stillinger, S. Torquato, Packing hyperspheres in high-dimensional Euclidean spaces. Phys. Rev. E 74, 041127 (2006).

74. S. Atkinson, F. H. Stillinger, S. Torquato, Detailed characterization of rattlers in exactly isostatic, strictly jammed sphere packings. Phys. Rev. E Stat. Nonlin. Soft Matter Phys. 88, 062208 (2013).

75. K. Roeller, J. Blaschke, S. Herminghaus, J. Vollmer, Arrest of the flow of wet granular matter. J. Fluid Mech. 738, 407–422 (2014).

76. M. Hennes, J. Tailleur, G. Charron, A. Daerr, Active depinning of bacterial droplets: The collective surfing of Bacillus subtilis. Proc. Natl. Acad. Sci., 201703997 (2017).

77. O. Praud, H. L. Swinney, Fractal dimension and unscreened angles measured for radial viscous fingering. Phys. Rev. E 72, 011406 (2005).

78. B. L. James, J. Kret, J. E. Patrick, D. B. Kearns, R. Fall, Growing Bacillus subtilis tendrils sense and avoid each other. FEMS Microbiol. Lett. 298, 12–19 (2009).

79. Y. Wu, H. C. Berg, Water reservoir maintained by cell growth fuels the spreading of a bacterial swarm. Proc. Natl. Acad. Sci. 109, 4128–4133 (2012).

80. A. Finkelshtein, D. Roth, E. Ben Jacob, C. J. Ingham, Bacterial Swarms Recruit Cargo Bacteria To Pave the Way in Toxic Environments. mBio 6 (2015).

81. D. Lopez, E. Lauga, Dynamics of swimming bacteria at complex interfaces. Phys. Fluids 26, 071902 (2014).

82. B. Ilkanaiv, D. B. Kearns, G. Ariel, A. Be’er, Effect of Cell Aspect Ratio on Swarming Bacteria. Phys. Rev. Lett. 118, 158002 (2017).

83. E. S. Gloag, et al., Micro-Patterned Surfaces That Exploit Stigmergy to Inhibit Biofilm Expansion. Front. Microbiol. 7 (2017).

84. M. Werb, et al., Surface topology affects wetting behavior of Bacillus subtilis biofilms. Npj Biofilms Microbiomes 3, 1–10 (2017).

85. D. A. Brock, T. E. Douglas, D. C. Queller, J. E. Strassmann, Primitive agriculture in a social amoeba. Nature 469, 393–396 (2011).

86. G. D. Palo, D. Yi, R. G. Endres, A critical-like collective state leads to long-range cell communication in Dictyostelium discoideum aggregation. PLOS Biol. 15, e1002602 (2017).

