## Supplemental Figures and Captions for "Physical factors contributing to regulation of bacterial surface motility"

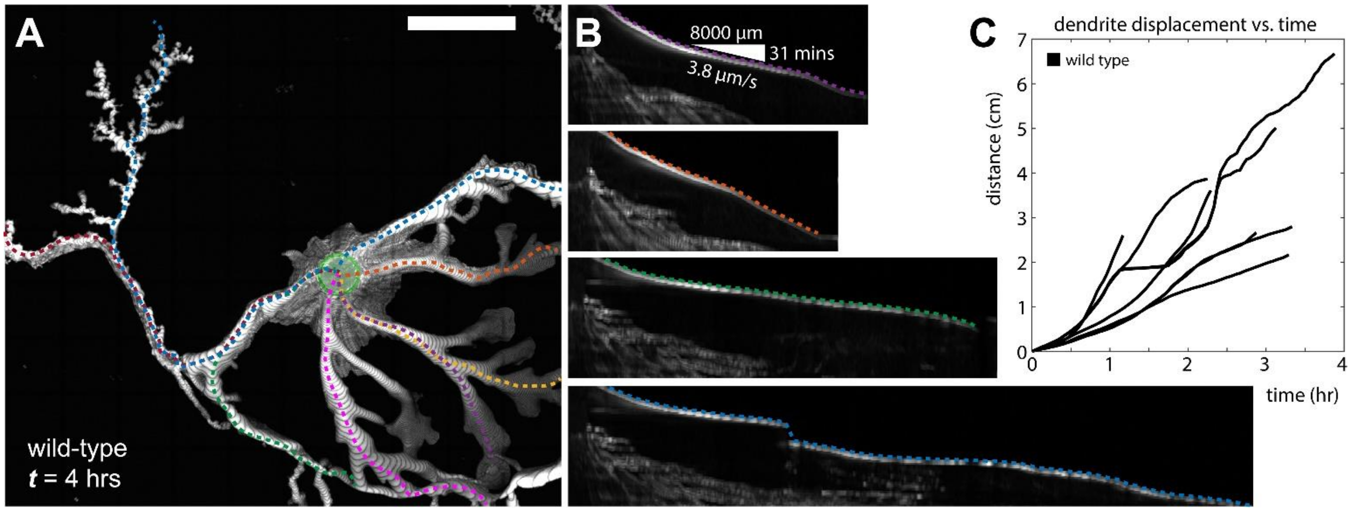


**SI Figure 1. Expansion dynamics of wild-type *Bacillus subtilis.*** **(A)** A maximum intensity projection of *B. subtilis* 3610 (wild type) spreading across nutrient-rich soft agarose (0.5% w/v). Colored contours (dashed lines) are drawn along selected dendrites. Original inoculation shown as a green circle. Scale bar is 1 cm. **(B)** Kymographs constructed from imaging data along specific contours (colors matched with (A)). The vertical dimension is time and the horizontal dimension is arc-length of the path of the dendrite. **(C)** Plot of dendrite displacement vs. time showing rapid surface motility ranging from ~4 µm/s up to ~10 µm/s. See Movie S2 for the complete time series.

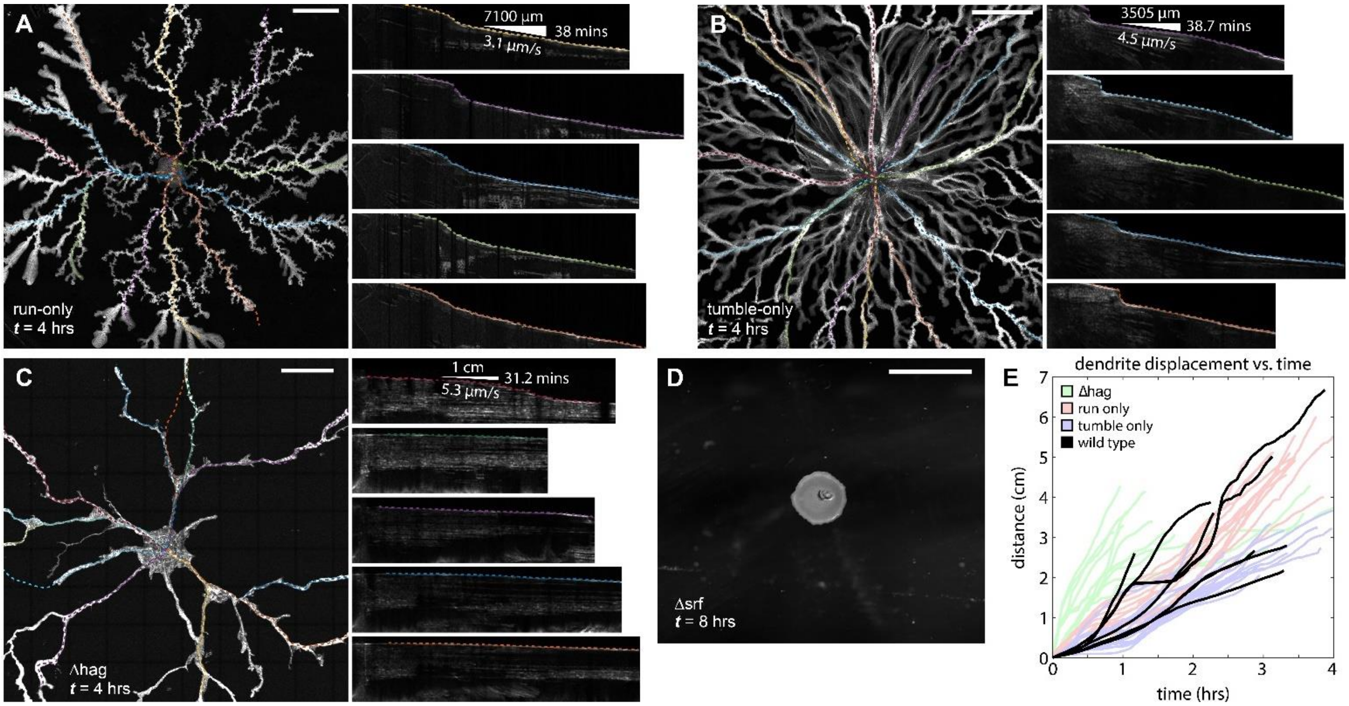


**SI Figure 2.** ***B. subtilis*** **genotypes that lack the ability to directly modulate their run-tumble frequency or are unable to self-propel still exhibit rapid dendritic expansion over surfaces.** Maximum intensity projections of dendrite dynamics for **(A)** run-only (Δ*CheB*), **(B)** tumble-only (Δ*CheY*), and **(C)** flagellar filament-deficient (Δ*hag*) genotypes, with contours (dashed lines) overlaid on dendrites and associated kymographs (right) with leading edges indicated (dashed lines). **(D)** Surfactin deficient (Δ*srf*) mutants do not exhibit dendritic expansion. **(E)** Each of the motility mutants (A-C) exhibit dendritic morphologies, with similar speeds for advancing dendrites as illustrated in this plot. Note that some dendrites of the Δ*hag* genotype – which cannot perform flagellar-mediated motility – outpace all other motile genotypes. These samples represent typical variation of advancing dendrites for their respective genotypes. All scale bars are 600 µm. See Movies S1, S3, S4, and S5.

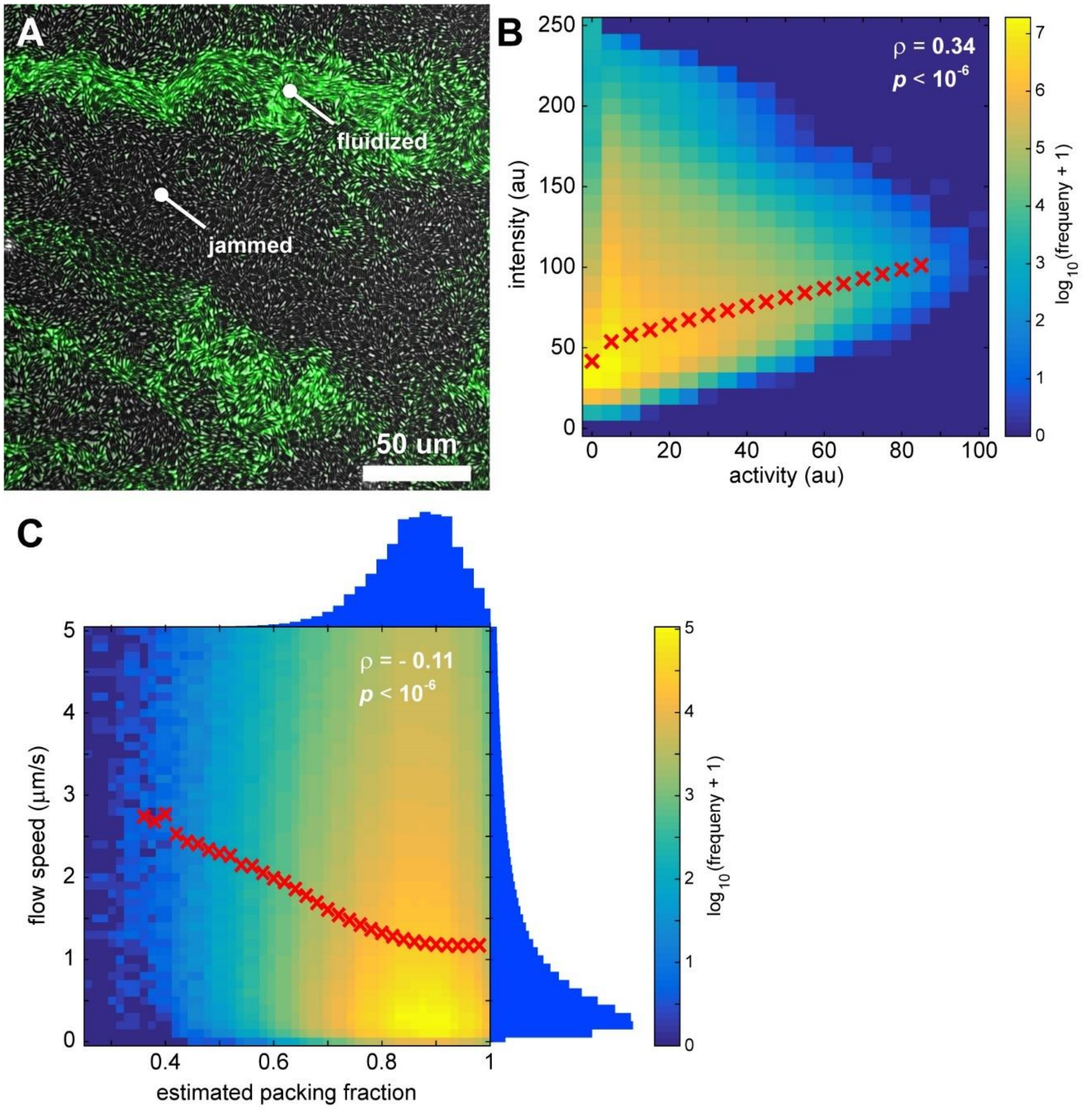


**SI Figure 3. Motion correlates with cell density. (A)** A snapshot from Movie S7C showing transiently fluidized cells (green, activity overlay) and jammed immobilized cells (gray). **(B)** Across 3000 frames, captured at 22 fps, we measured average intensity and average activity in blocks ~0.8 μm$^2$, and created a 2D histogram of those spatially correlated values (color map). For each column of intensity values at a fixed activity, we plot the mean intensity value (red X's) which shows a monotonic positive correlation, and calculated a bivariate correlation coefficient of 0.34, $p < 10^{-6}$. **(C)** Using particle-image velocimetry we measured the local flow speed of cells, and using an intensity threshold we measured the approximate packing fraction from the original images, both over blocks 12.4 μm$^2$. Those measures were significantly negatively correlated (-0.11, $p < 10^{-6}$), consistent with packing fraction impacting flow speed. The blue side histograms are the uncorrelated histograms for each measure. Note that in (B) and (C), the color scale is in powers of 10. The standard deviation for the null-hypothesis distribution in (B) was 7.2 x $10^{-5}$, and the standard deviation for the null-hypothesis distribution in (C) was 3.1 x $10^{-4}$.

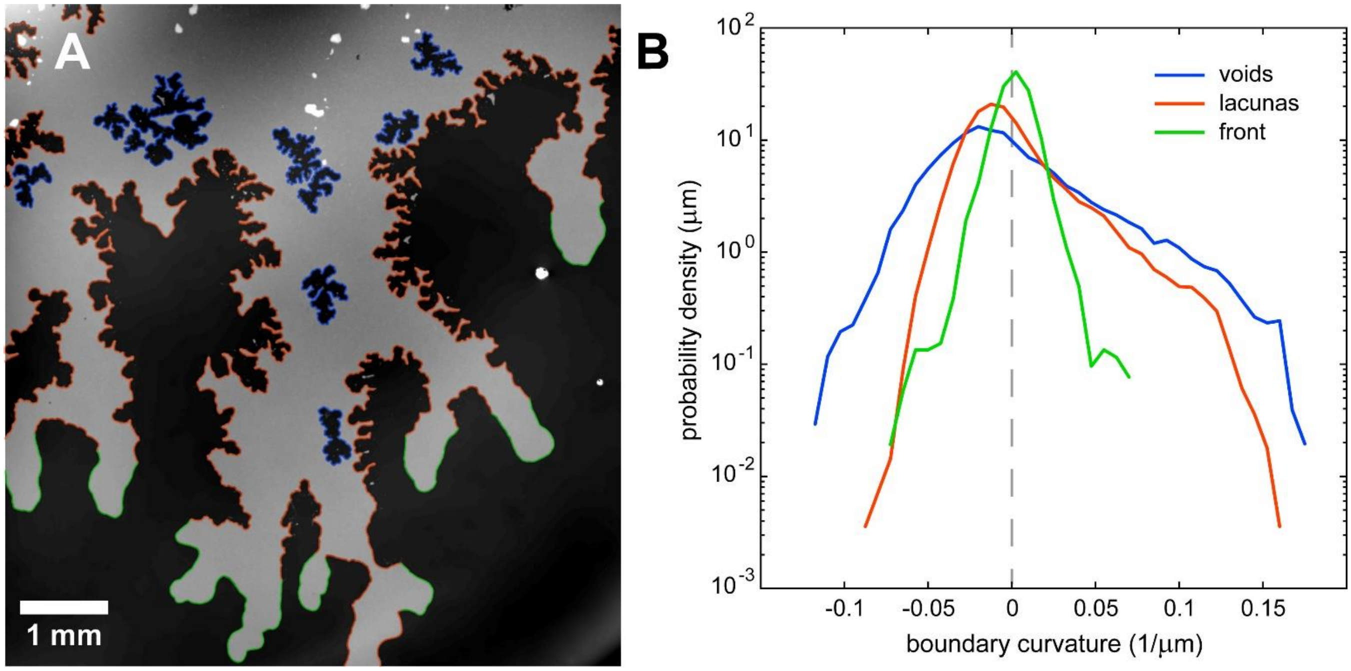


**SI Figure 4. Distributions of boundary curvature are consistent with outward pressure at the front. (A)** The underlying image is the same as Fig. 3A, showing the height of expanding dendrites on an agarose surface. Publically available image segmentation software [1] was used to identify the boundaries between agarose and cellular monolayers. Here void boundaries are colored blue, lacunar boundaries are colored orange, and boundaries of the advancing front are colored green. **(B)** We measured the distribution of curvatures for each type of boundary. Void and lacunar boundaries had similar distributions of curvature, both varying significantly more than curvatures at the advancing front, and both having negative modes consistent with negative pressure (and boundary retraction) in those regions. The front boundaries were significantly smoother (lower variation in curvature) and had a positive mode, consistent with positive pressure at the front.

[1] Ursell et al., Rapid, precise quantification of bacterial cellular dimensions across a genomic-scale knockout library (2017) *BMC Biology* 15:17.

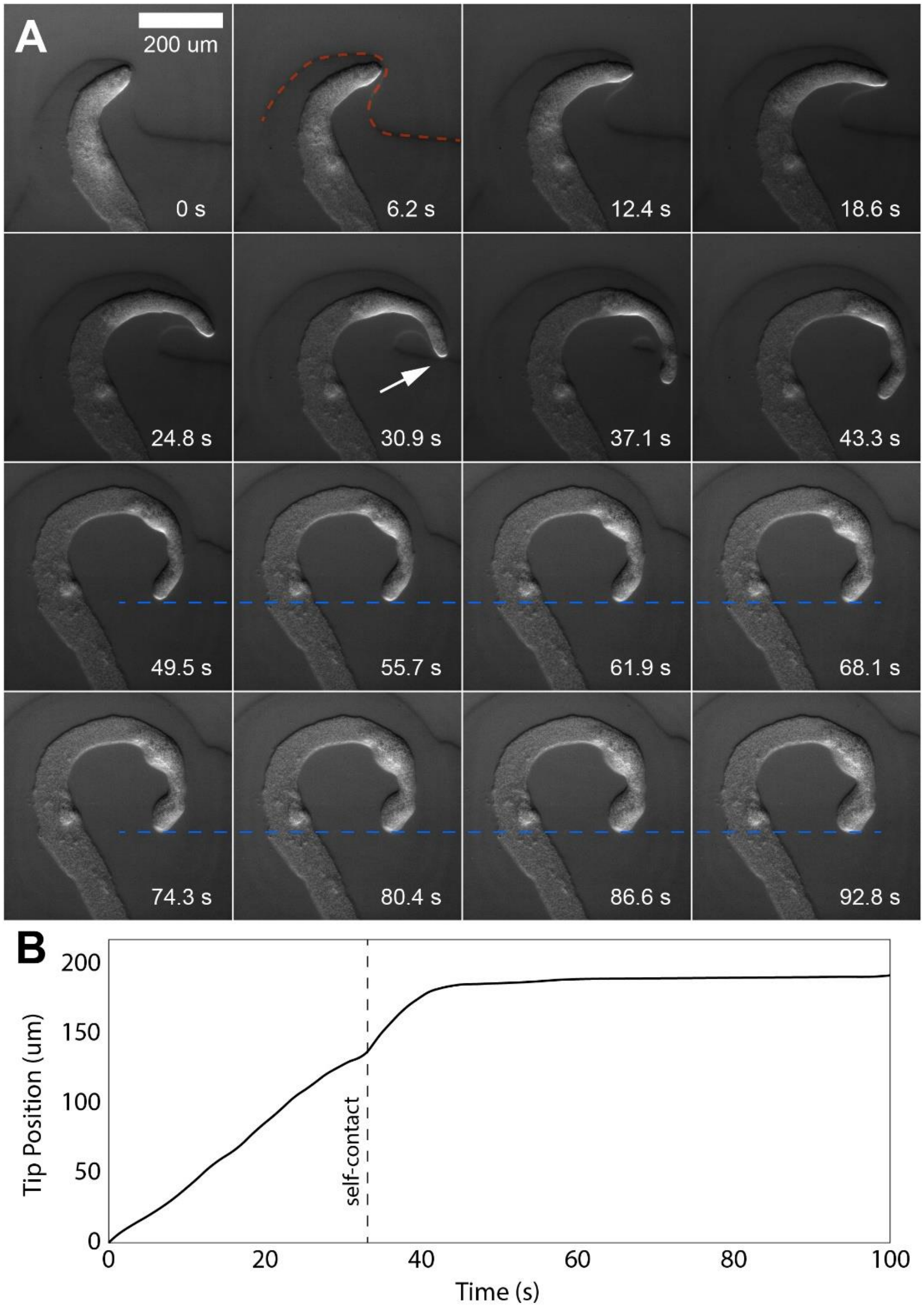


**SI Figure 5. Self-contact halts motion. (A)** Montage (and corresponding Movie S11) show the time-lapsed expansion of a dendrite with its accompanying wetting front imaged in oblique illumination. As a visual aid, the wetting front is outlined in the second pane (orange dashed line). The path of this dendrite curved and eventually contacted its own surfactin field (just after the white arrow). After self-contact, dendrite motion halted (blue dashed line) while cells within the dendrite tip remained motile and the number of cells in the tip increased as can be seen in the last eight panes. **(B)** Plot of the position of the advancing dendrite tip as a function of time. Up to the point of self-contact dendrite speed was steady, however dendrite motion slowed and halted ~15 s after self-contact, too quick to be related to changes in proliferation or phenotype.

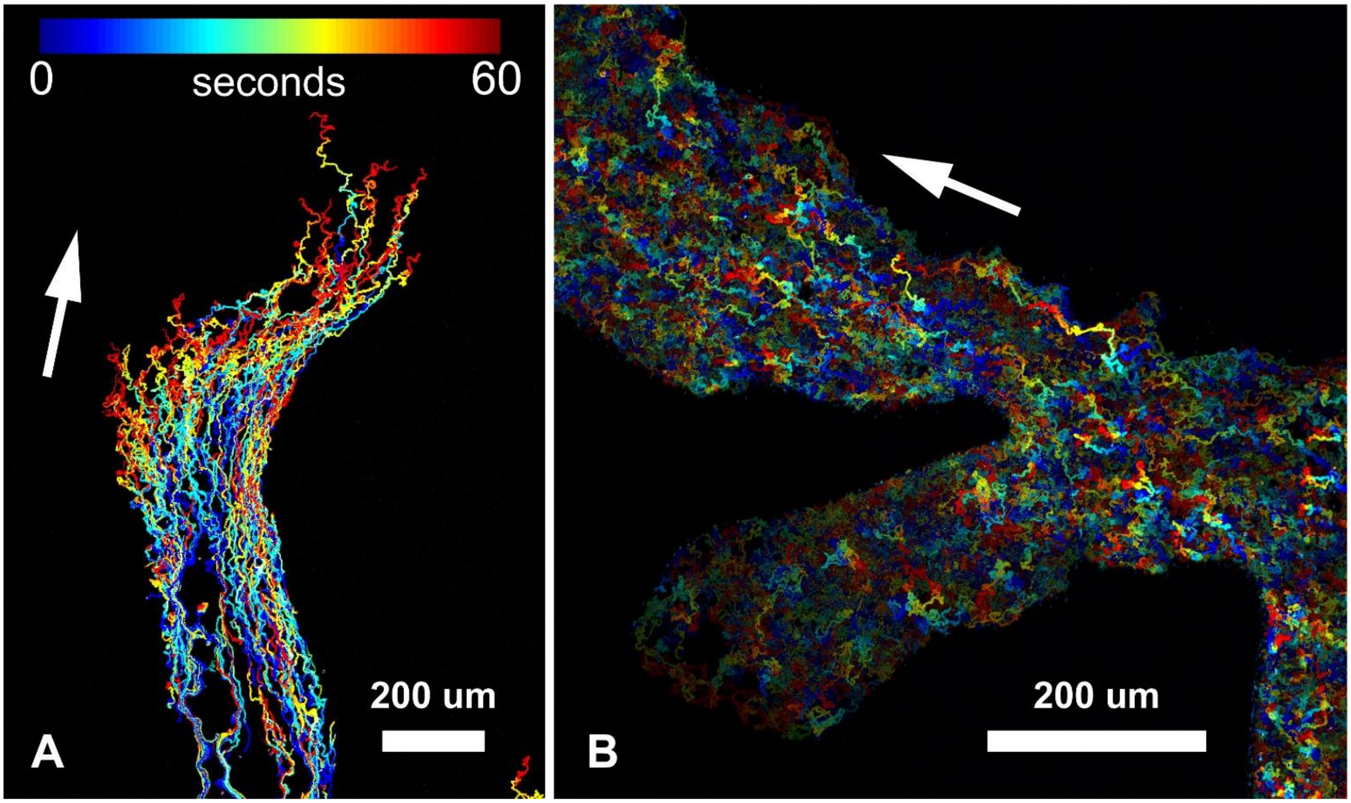


**SI Figure 6. Time-lapse bead movement in wild-type *B. subtilis*. (A)** Movement of 1 µm fluorescent tracer beads through time in a dendrite where net drift (flow speed) was ~ 2 µm/s. Note the erratic trajectories due to swarming of the surrounding cells. **(B)** Movement of 1 µm fluorescent tracer beads through time in a dendrite where net drift (flow speed) was < 1 µm/s. Note the erratic bead trajectories due to swarming of the surrounding cells, which prohibited usage of particle image velocimetry. In (A) and (B) the white arrows indicate the general direction of flow. See also Movies S14 and S15.

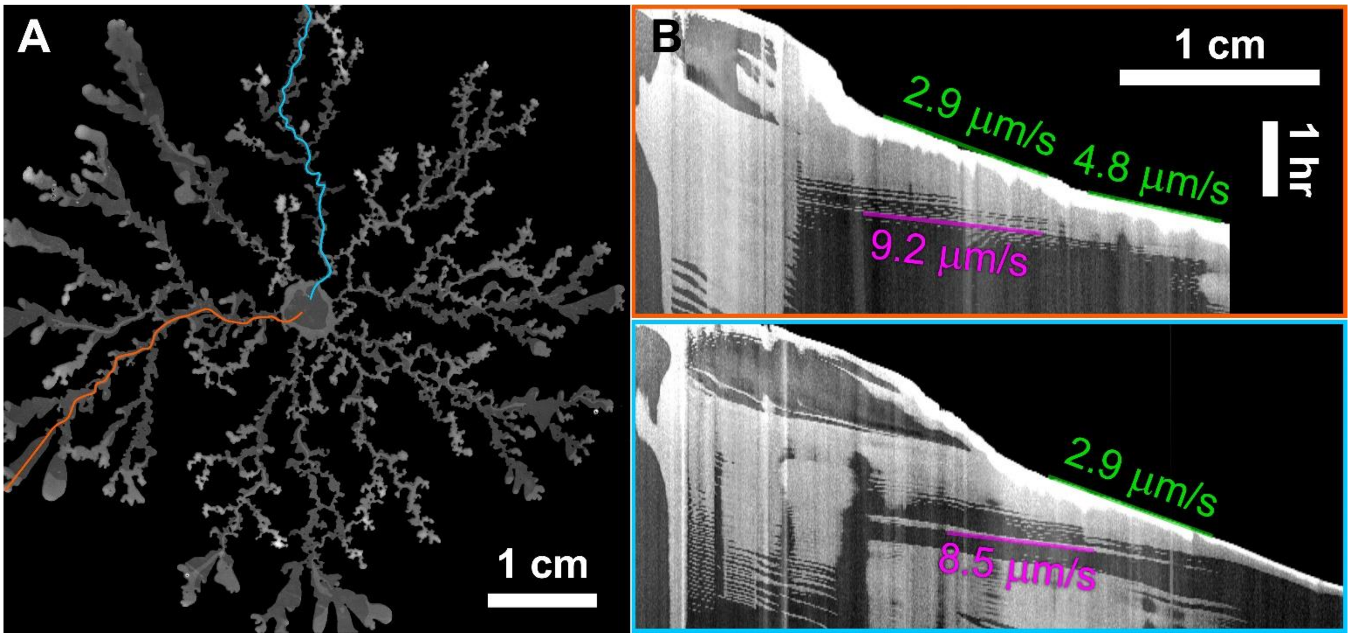


**SI Figure 7. Flow at tips vs. within a dendrite move at different speeds. (A)** A snapshot of the dendritic pattern of run-only *B. subtilis* covering an entire 10 cm plate. **(B)** Kymographs were constructed along the colored paths in the image. The tips extended at speeds of ~ 2 – 5 µm /s (green text) whereas internal flow, that is, flow originating near the parent colony moving along the dendrite outward to the tip, moved at significantly higher speeds as shown in (i) the overall kymographs, (ii) the specific measured speeds (magenta text), and (iii) along many dendrites in Movie S10. These data are consistent with the tracer bead data and with our conclusion that material can be transported from parent colony to the extending tip – specifically, due to the differential speed between tip and intra-dendrite flow seen here.
